# Phosphorylation of Claspin by elF2α kinase protects cells from heat stress

**DOI:** 10.1101/2025.08.21.670878

**Authors:** Chi-Chun Yang, Hidemasa Goto, Seiichi Oyadomari, Masai Hisao

## Abstract

Various biological stresses can induce replication stress, potentially leading to cancer development. Replication checkpoint protects cells against the threat of DNA damages by suspending cell cycle, allowing DNA repair. Chk1 kinase is activated by various stresses and is required for replication checkpoint. An adaptor protein, Claspin, mediates Chk1 activation, but specific mechanisms involved are still unknown. We demonstrate that heat stress induces hyperphosphorylation of Claspin, resulting in Chk1 activation. Claspin phosphorylation is impaired in cells lacking two eIF2α kinases, GCN2 and HRI, which are parts of the integrated stress response (ISR) pathway. Mass spectrometry identified five heat-inducible phosphorylation sites in the C-terminal region of Claspin, and Claspin lacking these sites failed to activate checkpoint. Our data support a model that the unphosphorylated C-terminal tail of Claspin masks its CKBD (Chk1 Binding Domain)-AP (Acidic Patch) domain, preventing it from interacting with Cdc7. Phosphorylation of the tail causes it to dissociate from CKBD-AP and facilitates Cdc7 interaction, leading to increased Chk1 activation. Our findings demonstrate crosstalk between replication checkpoint and ISR pathways, and suggest potential new strategies for cancer therapy.

## Introduction

The faithful duplication of the genome is crucial for maintaining cellular homeostasis and preventing diseases such as cancers. The process of DNA replication is constantly challenged by various endogenous and exogenous obstacles that can impede the progression of replication forks, leading to a phenomenon known as replication stress. To safeguard genomic integrity, cells have evolved intricate mechanisms, collectively referred to as the DNA replication checkpoint, to detect and respond to replication stress. The ATR (Ataxia Telangiectasia and Rad3-related)-Chk1(Checkpoint Kinase 1) signaling pathway is central to this checkpoint response, stabilizing stalled replication forks and coordinating DNA repair processes. Claspin, a key mediator protein, facilitates the activation of Chk1 by ATR in response to replication stress (1–4). In addition, Cdc7 or CK1γ1 can phosphorylate CKBD (Chk1 Binding Domain) of Claspin to induce Chk1 activation (5, 6). In yeast, the Claspin homolog, Mrc1, also plays a crucial role in DNA replication checkpoint, transmitting the replication stress signal to downstream effector kinases like Rad53 and Cds1(7–9). Beyond its well-established role in the replication stress response, Claspin/Mrc1 has important functions in both the initiation and elongation phases of DNA replication (10–12), ensuring the timely and accurate duplication of the genome. It also plays a role in histone distribution and epigenetic regulation (13,14). The abundance of Claspin is regulated during the cell cycle through ubiquitination-proteasome-mediated degradation, regulating replication checkpoint signaling (15).

It was reported that Mrc1 in budding yeast is targeted by multiple stress-activated protein kinases (SAPKs) including Hog1, Mpk1, and Psk1, which are activated by osmotic stress, heat stress, and oxidative stress, respectively. These SAPKs phosphorylate Mrc1 at its N-terminal region, leading to a delay of the cell cycle, preventing transcription-associated recombination (TAR) and genomic instability, and maximizing cell viability. The N-terminal phosphorylation of Mrc1 acts as a key integrator of multiple signals to block DNA replication and prevent transcription-replication conflicts, ensuring genomic integrity during transcriptional outbursts or unscheduled transcription during S phase (16,17).

In mammalian cells, we and others previously reported that Chk1 can be activated by various biological stresses, including hypoxia, osmotic shock, oxidative stress, LPS (bacterial infection), nutrient deprivation, ER stress, unfolded protein response, and heat, in a manner dependent on Claspin (19–26). It has been reported that Claspin is directly phosphorylated by p38 MAP kinase, the mammalian homolog of Hog1 kinase, which facilitates the repair of lesions and protects cells from DNA damage in response to osmotic stress during S-phase. However, mechanisms of Chk1 activation by other biological stresses, potential crosstalk between other stress response pathways and replication checkpoint, and the role of Claspin in these processes are largely unknown in higher eukaryotes.

Besides DNA replication checkpoint, cells possess other adaptive mechanisms to cope with stressors. The heat-shock response (HSR) is a highly conserved cellular defense mechanism activated by stressors, including heat, oxidative stress, and toxins (27). This response centers around the induction of heat shock proteins (HSPs), which function as molecular chaperones to maintain proteostasis by aiding in protein folding, preventing aggregation, and assisting in protein repair or degradation (28–30). HSPs, including HSP100, HSP90, HSP70, and smaller HSPs, play crucial roles in cellular homeostasis (31–33) and stress management, and are implicated in human diseases such as cancers and neurodegenerative disorders (34). Heat shock factors (HSFs), particularly HSF1, orchestrate the transcriptional activation of HSPs during stress (35). The interplay between HSPs and cellular pathways involved in apoptosis, immune responses, and inflammation, highlights their multifunctional roles in cellular adaptation and survival (36,37). The heat shock response also has therapeutic potential, with ongoing research targeting HSPs and their co-chaperones exploring applications for treating various diseases (38).

Several cellular stresses are known to activate the integrated stress response (ISR) in which the eukaryotic translation initiation factor 2 (eIF2α) is phosphorylated at serine-51, to inhibit translation (39). Four eIF2α kinases, heme-regulated inhibitor (HRI), protein kinase R (PKR), PKR-like endoplasmic reticulum kinase (PERK), and general control non-depressible 2 (GCN2) are activated by heme depletion, viral infection, endoplasmic reticulum stress, and amino acid starvation, respectively (40, 41). In this report, we discovered that heat-induced checkpoint activation also depends on ISR, and eIF2α kinases play a role in this process by phosphorylating Claspin. These results demonstrate crosstalk between the replication checkpoint and ISR pathways.

## Results

### Phosphorylation of residues near the C-terminus of Claspin is Critical for Checkpoint Activation and Cell Survival Upon Heat Stress

We previously reported that Claspin undergoes a mobility shift on SDS-PAGE under heat shock stress. U2OS cells were synchronized by arresting them at G2/M phase using nocodazole and then releasing them at either 37°C or 42°C. In control 37°C cultures, a relatively faint mobility shift was observed at 3-6 hrs after release, indicating a cell cycle-dependent phosphorylation of claspin (**Fig. 1A**, lanes 8 and 9). In cells exposed to 42°C heat stress, an additional intense shift was observed between 12 and 15 hrs after release, corresponding to S-phase as determined by the Cdt1 and Geminin expression (**Fig. 1A**, lanes 5 and 6). Chk1 activation followed a pattern very similar to Claspin phosphorylation. As expected, Claspin phosphorylation and Chk1 activation were not observed in Claspin-depleted cells even after heat treatment (**Fig. 1A**, lanes 14, 15, 17 and 18). We next analyzed the DNA content of cells using FACS (**Fig. 1B**). In control (37°C) and heat-stressed (42°C) cells, cells transited from 4C to 2C within 3 hrs after release. Control cells started to enter S phase at 9 hrs after release in both the presence of Claspin (siControl) and when Claspin was depleted (siClaspin). In heat-stressed cells, the transition to the 2C state was not affected but progression into S phase was inhibited. In heat-treated siClaspin conditions, cells with 2C DNA content decreased with concomitant increase of subG1 DNA populations (representing dead cells) at 12-15 hrs after release, probably representing cell death after aberrant mitosis (**Fig. 1B**).

**Figure 1.**
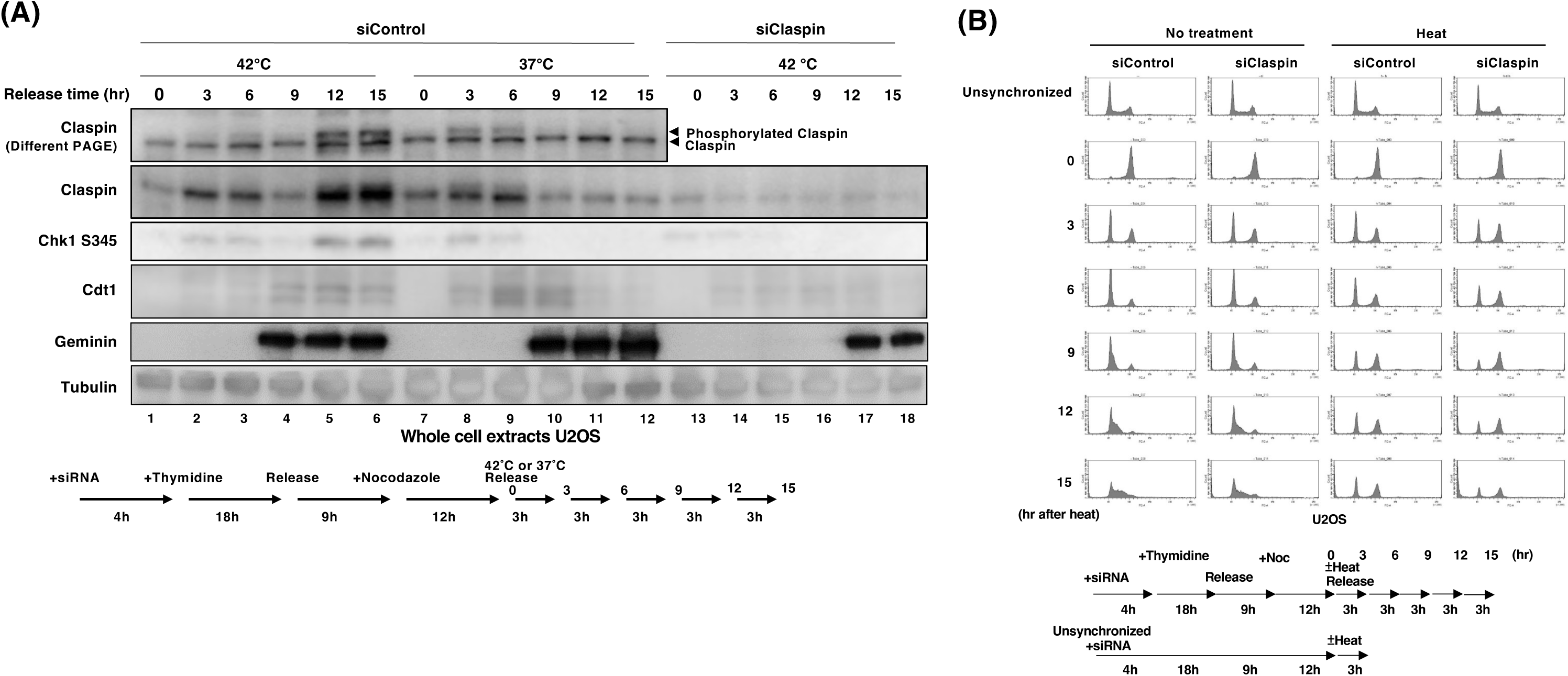
Claspin-dependent Chk1 activation during heat treatment. U2OS cells were treated with control siRNA (siControl) or Claspin siRNA (siClaspin) and synchronized at the G2/M phase using a thymidine block followed by nocodazole treatment, as indicated in the scheme presented under the panels. After release from the nocodazole block, cells were cultured at 37°C or 42°C and harvested at the indicated time points. (**A**) The whole-cell extracts were analyzed by western blotting with the indicated antibodies. (**B**) Cell cycle analysis of the cells stained with propidium iodide (PI) by flow cytometry.

To further explore heat-stress-induced changes, we analyzed Claspin phosphorylation sites in 293T cells cultured for 2 hrs at 37°C or 42°C. Five novel phosphorylated amino acids (T1322, S1325, T1326, S1327, and T1330) were identified near the C-terminus of Claspin after 42°C treatment (**Supplementary Fig. S1A**). Moreover, we identified a potential phosphorylation motif between aa1321–1331 (**T**DD**STS**GL**T**RS) of human Claspin, which is conserved in mice, cows, and frogs (**Supplementary Fig. S1B**). A similar potential phosphorylation site was not detected in fission yeast Mrc1.

To evaluate the biological significance of these five phosphorylation sites, we generated two types of mutants, ST/A, in which all serines (S) and threonines (T) were mutated to alanine (A), and ST/E, in which they were mutated to glutamic acid (E). We then established stable clones expressing either wild-type or mutant Claspin in HCT116 Claspin-AID-myc cells. By adding auxin to induce the degradation of endogenous Claspin, we can assess the effects of these exogenous Claspin variants. First, we evaluated whether these five phosphorylation sites regulate checkpoint activation upon heat shock stress. Chk1 was phosphorylated under heat-stressed conditions in the presence of endogenous Claspin (**Fig. 2A**, lanes 9 and 10; **Fig. 1 and Supplementary Fig. S2**), and this phosphorylation was abolished upon Claspin depletion. Chk1 phosphorylation was restored when wild-type or ST/E mutant Claspin was expressed, but not when the ST/A mutant was expressed (**Fig. 2A**, lanes 12, 14, and 16). Furthermore, wild-type Claspin showed a noticeable mobility shift upon heat stress, whereas the ST/A mutant did not (**Fig. 2B**). These results indicate that phosphorylation at the identified C-terminal amino acids of Claspin is crucial for heat stress-dependent checkpoint activation. We next examined whether mutations at these sites affect cell survival in heat-stressed conditions. To investigate this, we used colony formation assays to assess cell survival rate at 42°C. At the 0 min time point, there was a slight decrease in cell survival in the absence of Claspin, but no significant difference in survival when wild-type, ST/A, and ST/E mutants were expressed. However, after 90 min at 42°C, the number of colonies in Claspin-deficient cells (+auxin) decreased by 44% compared to the control (-auxin). Expression of wild-type Claspin or the ST/E mutant restored the colony formation, but expression of the ST/A mutant (37% decrease) did not (**Fig. 2C**). These findings suggest that preventing C-terminal phosphorylation of Claspin results in a deficiency in checkpoint activation and increased cell death upon heat shock stress.

**Figure 2.**
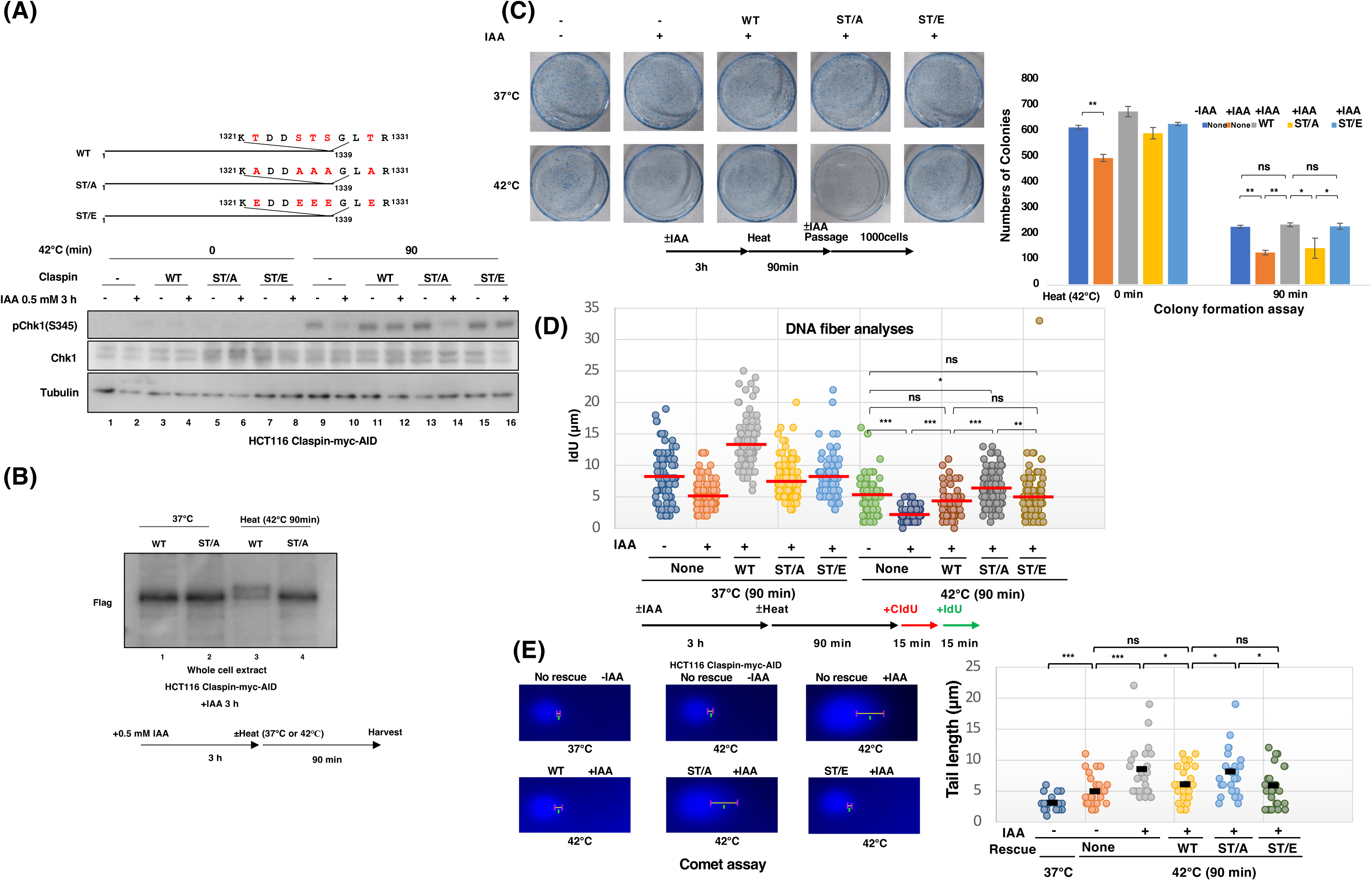
Role of Claspin C-terminal phosphorylation in checkpoint activation and cell survival under heat shock stress. (**A**) Western blot analysis of HCT116 Claspin-myc-AID cells stably expressing either wild-type Claspin-Flag (WT), ST/A-Claspin-Flag (ST/A), or ST/E-Claspin-Flag (ST/E) were treated with or without 0.5 mM auxin for 3 hrs. Cells were then cultured at 42°C for 0 or 90 min. Whole cell extracts were analyzed using the indicated antibodies. Sequences of the wild-type and the mutant are shown. (**B**) HCT116 Claspin-myc-AID cells, stably expressing either WT or ST/A, were treated with 0.5 mM auxin for 3 hrs. Cells were then cultured at 37°C or 42°C for 90 min. Whole cell extracts were analyzed using the DDDK antibody. (**C**) Colony formation assay of Claspin mutant cell lines under heat stress conditions. HCT116 Claspin-myc-AID cells stably expressing either WT, ST/A, or ST/E Claspin were treated with or without 0.5 mM auxin for 3 hrs and cultured at 37°C or 42°C for 90 min. Cells were plated at a density of 1,000 cells per dish in medium with or without auxin. After 3 weeks, colonies were stained with Coomassie Brilliant Blue (CBB) and counted. Statistical significance is indicated as follows: ns (p > 0.05); * (p ≤ 0.05); ** (p ≤ 0.01); *** (p ≤ 0.001). (**D**) DNA replication analysis using the DNA fiber assay. HCT116 Claspin-myc-AID cells, stably expressing either WT, ST/A, or ST/E Claspin, were treated with or without 0.5 mM auxin for 3 hrs and cultured at 37°C or 42°C for 90 min. Cells were pulse-labeled with CldU and IdU. DNA fibers were stained with anti-CldU and anti-IdU antibodies, followed by Alexa-555 and Alexa-488 secondary antibodies. The slides were mounted and observed under a fluorescence microscope. The length of IdU fibers was measured, with red lines indicating average values. (**E**) Comet assays of HCT116 Claspin-myc-AID cells, stably expressing either WT, ST/A, or ST/E Claspin, treated with or without 0.5 mM auxin for 3 hrs and cultured at 37°C or 42°C for 90 min. After electrophoresis, the cells were stained with DAPI and observed under a fluorescence microscope. Tail length was measured, with the scale indicating its length. The tail length of each cell is shown in the graph. Statistical significance is indicated as follows: ns (p > 0.05); * (p ≤ 0.05); ** (p ≤ 0.01); *** (p ≤ 0.001).

We next sought to understand how phosphorylation at these five amino acids promotes cell survival under heat shock stress. First, we investigated replication speed using a DNA fiber assay. DNA was labeled with CldU and IdU consecutively for 15 min each, and the IdU length of the CldU-IdU stretch was measured to assess the speed of the moving fork. We observed a decrease in replication speed by 38% in Claspin-deficient cells, consistent with previous reports demonstrating that Claspin facilitates replication fork progression under normal growth conditions (10–12). At 42°C, fork rate was reduced in the wild-type cells, and was further reduced by 50% after depletion of Claspin. While expression of the wild-type Claspin or the ST/E mutant restored fork speed to the heat-treated wild-type level, expression of the ST/A mutant increased speed to the level close to that of the non-treated wild-type (**Fig. 2D**). These results suggest two functions of Claspin in regulating replication fork speed. First, Claspin is important for normal fork progression. Second, phosphorylation of C-terminal sites on Claspin inhibits fork speed, possibly through checkpoint activation.

Next, we examined whether heat stress leads to increased DNA damage using comet assays. In heat stressed conditions, Claspin depletion increased comet tail length by 40%. This increase in length did not occur upon expression of the wild-type or ST/E Claspin, but did occur upon expression of the ST/A mutant (38% increase; **Fig. 2E**). Altogether, these results indicate that C-terminal phosphorylation of Claspin plays a crucial role in checkpoint activation, and preventing phosphorylation increases DNA damage and render cells more sensitive to heat stress.

### HRI and GCN2 Kinases Mediate Claspin Phosphorylation and Checkpoint Activation Upon Heat Stress

Claspin has been reported to be phosphorylated by Cdc7 and CK1γ1 upon replication stress (5,6), by PI3K following release from serum starvation (42), and by p38 during osmotic stress (21). Similarly, the fission yeast Mrc1 has been shown to be phosphorylated by Mpk1 under heat shock stress (17). To identify kinases that phosphorylate mammalian Claspin during heat shock stress, we analyzed the sequences surrounding the C-terminal phosphorylation sites (T1322, S1325, T1326, S1327, and T1330) using the Kinase Library (43,44) (**Fig. 3A**). Results predicted that S1325 and T1326 are highly likely to be phosphorylated by HRI, GCN2, or PERK, which are eIF2α kinases known to mediate integrated stress responses (41). Notably, HRI kinase has been reported to be activated upon heat shock stress in association with Hsp90 (45). To examine whether the eIF2α kinases are required for heat-dependent Chk1 activation, we used MEF cells in which all the four eIF2α kinases are knocked out (4KO) and 4KO cells expressing each kinase separately (4KO + PERK, 4KO + PRK, 4KO + HRI, and 4KO + GCN2). Chk1 phosphorylation was absent in 4KO cells cultured at 42°C, and restored in 4KO + HRI and 4KO + GCN2 MEF cells but not in 4KO + PERK or 4KO + PRK cells (**Fig. 3B**). Furthermore, when HRI or GCN2 were knocked down by siRNA in 4KO + HRI and 4KO + GCN2 MEF cells, under heat stress conditions, Chk1 phosphorylation as well as RPA phosphorylation (an indicator for replication stress/ DNA damage) was reduced (**Fig. 3C**). These findings indicate that HRI and GCN2 play critical roles in replication checkpoint activation during heat shock stress.

**Figure 3.**
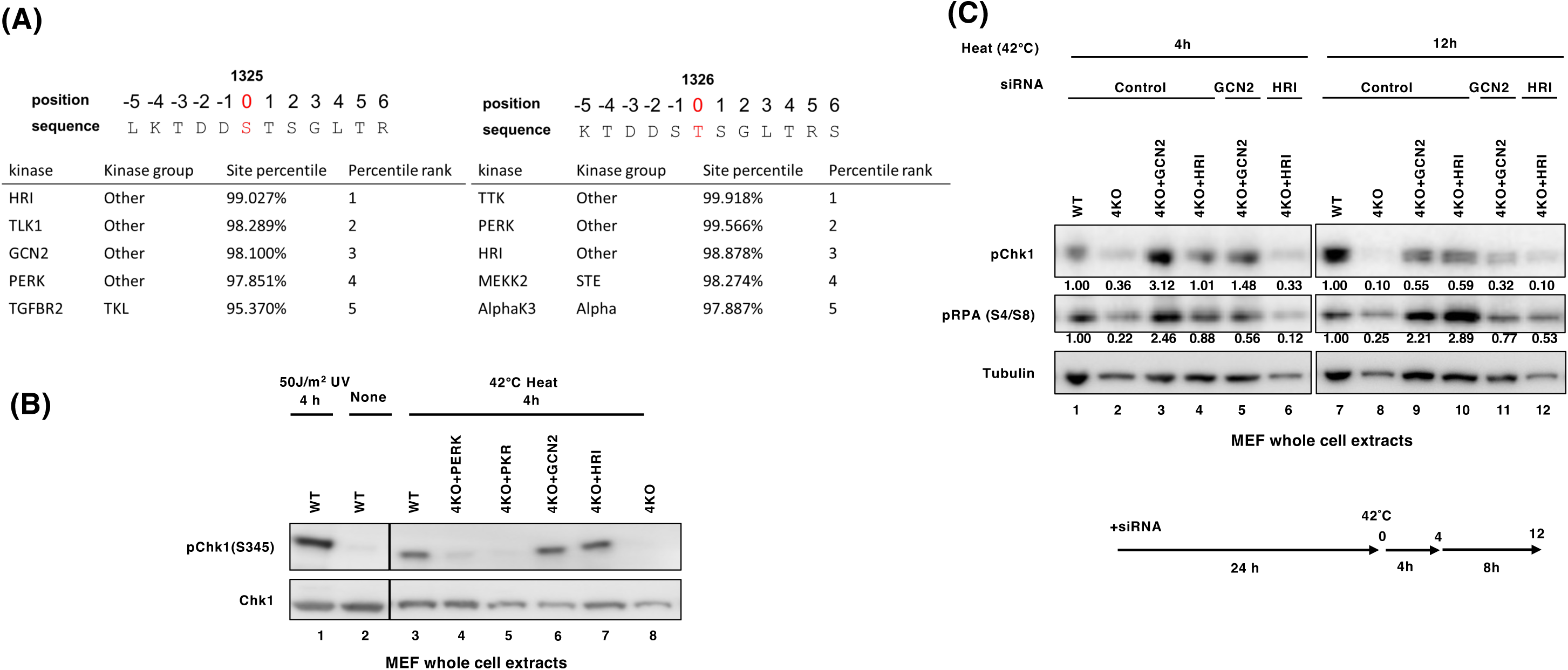
HRI and GCN2 regulate checkpoint activation under heat shock stress. (**A**) Kinase prediction for the C-terminal phosphorylation sites (right, S1325; left, T1326) of Claspin under heat shock stress. (**B**) Wild-type, 4KO (lacking PERK, PKR, HRI, and GCN2), 4KO + PERK(stably expressed), 4KO + PKR, 4KO + HRI, and 4KO + GCN2 MEF cells were cultured at 42°C for 4 hrs. Whole cell extracts were harvested and analyzed by Western blotting with the indicated antibodies. (**C**) The same set of cells as in **B** were transfected with or without siHRI or siGCN2 for 24 hrs, and then cultured at 42°C for 4 or 12 hrs, and were harvested. Whole cell extracts were analyzed by Western blotting with the indicated antibodies.

### HRI and GCN2 Directly Phosphorylate Claspin and are Required for Cell Viability Under Heat Stress

To examine whether HRI or GCN2 directly phosphorylates the C-terminus of Claspin, we conducted kinase assays using purified proteins and found that both kinases could phosphorylate wild-type Claspin (**Fig. 4A**, lanes 4 and 7). Furthermore, HRI and GCN2 exhibited significantly lower phosphorylation activity on the ST/A mutant suggesting that these kinases phosphorylate the identified C-terminal Claspin sites (**Fig. 4A**, lane 5). Interestingly, the ST/E mutant was phosphorylated even in the absence of an added kinase, suggesting the presence of a kinase copurifying with this mutant (see below).

**Figure 4.**
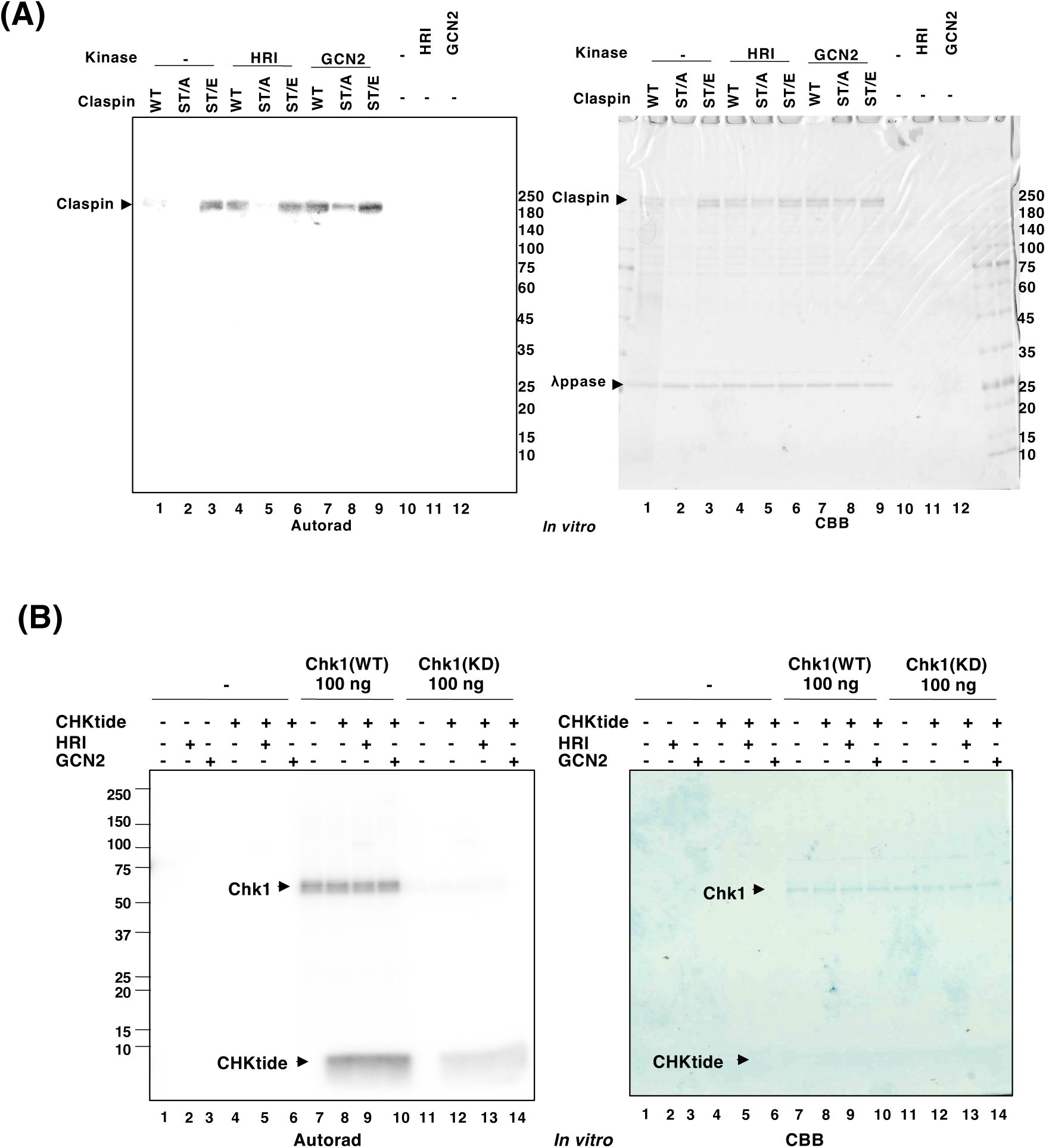
HRI and GCN2 phosphorylate Claspin *in vitro*. (**A**) Purified wild-type, ST/A, and ST/E mutant Claspin proteins were treated with λ phosphatase (λPPase), pre-treated with PhosSTOP™, and then mixed with purified HRI or GCN2 (20 ng each) and incubated in kinase buffer at 30°C for 30 min. (**B**) Wild-type Chk1 (WT) or a kinase-dead Chk1 mutant (D130A, KD) protein was mixed with HRI or GCN2 kinase (20 ng each) and incubated in kinase buffer with CHKtide polypeptide at 30°C for 60 min. In (**A**) and (**B**), the reaction mixtures were subjected to SDS-PAGE. The gel was stained with Coomassie Brilliant Blue (CBB) (right panel) and then autoradiographed (left panel).

Further supporting the idea that GCN2 phosphorylates Claspin, Claspin expressed in heat-stressed HCT116 cells migrates as a doublet on polyacrylamide gels (**Fig. 5A**).

**Figure 5.**
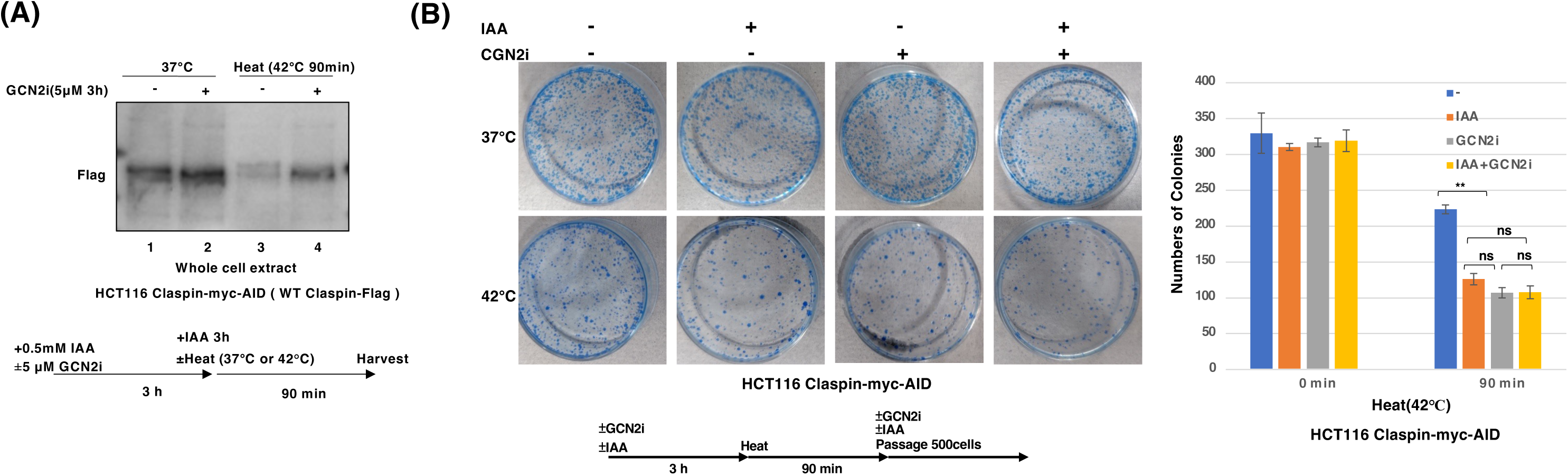
GCN2 and Claspin collaborate to regulate cell viability under heat shock stress. (**A**) HCT116 Claspin-myc-AID cells stably expressing wild-type Claspin-Flag were treated with auxin and with or without GCN2 inhibitor (GCN2-IN-1, A-92) for 3 hrs. The cells were then cultured at 37°C or 42°C for 90 min. Whole cell extracts were harvested and analyzed by Western blotting using the anti-DDDK antibody. (**B**) HCT116 Claspin-myc-AID cells, treated with or without 0.5 mM auxin and 10 nM GCN2 inhibitor for 3 hrs, were cultured at 37°C or 42°C for 90 min. Cells were plated at a density of 500 cells per dish in medium with or without auxin. After 3 weeks, colonies were stained with Coomassie Brilliant Blue (CBB) (left) and counted (right). Statistical significance is indicated as follows: ns (p > 0.05); * (p ≤ 0.05); ** (p ≤ 0.01); *** (p ≤ 0.001).

However, in cells treated with a GCN2 inhibitor (GCN2i), the upper band of the doublet disappears.

To determine whether HRI and GCN2 can also directly phosphorylate Chk1, we again used purified proteins. As previously reported (46,47), Chk1 can phosphorylate itself as well as its substrate, CHKtide (**Fig. 4B**, lanes 7 and 8). However, Chk1 phosphorylation and kinase activity did not increase when mixed with HRI or GCN2 (**Fig. 4B**, lanes 8 and 9), suggesting that HRI and GCN2 do not directly phosphorylate Chk1.

We then assessed cell viability using colony formation assays. Under heat stress conditions, the cell survival rate of Claspin-deficient cells was 56% of that of the control cells, while the survival rate of cells treated with GCN2i was similar (48%). The survival rate of GCN2i-treated Claspin-deficient cells was 48%, a value similar to that of Claspin-depleted and GCN2i-treated cells (**Fig. 5B**). Thus, Claspin deficiency and GCN2 inhibition do not have additive effects, suggesting that eIF2α kinases and Claspin operate through the same pathway to activate checkpoint and regulate cell survival after heat shock.

### ATR mediates checkpoint activation during heat shock stress

ATR is a key protein kinase activated in response to replication stress (48). It initiates a signaling cascade that leads to cell cycle arrest, allowing time for resumption of DNA replication and DNA repair. Claspin acts as a mediator in this pathway and facilitates the activation of Chk1 by ATR. Chk1 then phosphorylates various downstream targets, including cell cycle regulators, to halt cell cycle progression. The ATR-Claspin-Chk1 axis is essential for maintaining genome stability and preventing the propagation of damaged DNA after replication stress.

We sought to elucidate whether ATR is involved in the regulation of checkpoint activation during heat shock stress. Auxin-inducible Claspin-myc-AID HCT116 cells expressing either wild-type Claspin or ST/A and ST/E mutants were treated with or without the ATR inhibitor, AZD6738, and subsequently incubated at 42°C for 90 min. Heat stress induced Chk1 phosphorylation when WT Claspin and ST/E mutants were expressed, but not when the ST/A mutant was expressed (**Fig. 6A**). Notably, the addition of the ATR inhibitor resulted in a reduction in Chk1 phosphorylation in cells expressing the wild-type and ST/E mutants, strongly suggesting that ATR contributes to Chk1 activation during heat stress. Supporting this idea, we detected physical interactions between ATR and Claspin by immunoprecipitation (**Supplementary Fig. S3**). This interaction occurred between ATR and wild-type Claspin and ST/E and ST/A mutants. Thus, while ATR is needed for heat stress-dependent Chk activation, physical Claspin-ATR interactions occur independently of C-terminal Claspin phosphorylation.

**Figure 6.**
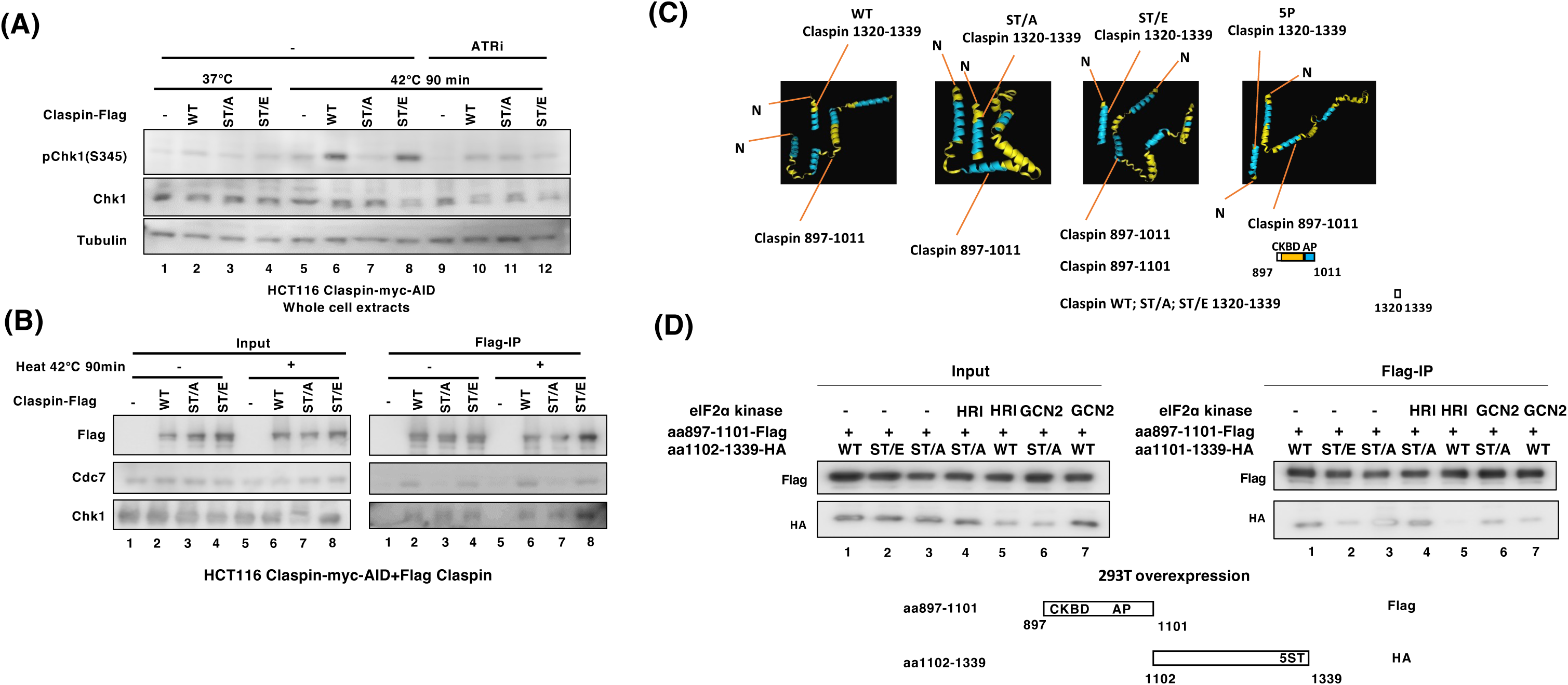
HRI or GCN2-mediated phosphorylation of the C-terminal segment inhibits interaction with the CKBD-AP segment, leading to Cdc7 recruitment to Claspin. (**A**) HCT116 Claspin-myc-AID cells, stably expressing wild-type (WT), ST/A, or ST/E Claspin, were treated with auxin in the presence and absence of ATR inhibitor (ATRi; AZD6738) for 3 hrs. Cells were then cultured at either 37°C or 42°C for 90 min. Whole-cell extracts were collected and analyzed via western blotting with the indicated antibodies. (**B**) The same cells used in (**A**) were treated with auxin, cultured at 37°C or 42°C for 90 min, and subjected to pull-down assays using M2 Flag beads. Co-immunoprecipitated proteins were analyzed via western blotting to detect the indicated proteins. (**C**) Predicted structural models of Claspin CKBD-AP polypeptide (aa 897–1011) in a complex with the Claspin C-terminal polypeptide (aa 1320–1339; WT, ST/A, ST/E and 5P) generated by AlphaFold3. 5P represents the peptide with 5 potential HIR/GCN2-mediated phosphorylation sites (serine and threonine) being phosphorylated. The WT and ST/A mutant could interact with the AP segment or could be accommodated into the pocket segment, whereas ST/E or the phosphorylated version of the C-terminal polypeptide is predicted not to interact with the pocket. (**D**) 293T cells were co-transfected with plasmids expressing HA-tagged Claspin C-terminal polypeptide (aa1101–1339, #28; WT, ST/A, or ST/E) and Flag-tagged Claspin AP polypeptide (aa 987–1100), as well as the plasmid expressing HRI or GCN2. Cell lysates were immunoprecipitated using anti-Flag (M2) beads, and the presence of the C-terminal polypeptide was examined.

### C-terminal phosphorylation regulates Cdc7 binding to Claspin

Cdc7, a serine/threonine kinase, plays a pivotal role in initiation of DNA replication by phosphorylating critical components of the pre-replicative complex (49, 50). CK1γ1, another serine/threonine kinase, also regulates various cellular processes, including DNA replication and repair (7,8). Previously, we and others reported that Cdc7 and CK1γ1 interact with and phosphorylate Claspin to facilitate Chk1 activation (6). Thus, we next examined the interaction between Claspin and Cdc7 and CK1γ1 upon heat stress. Whereas CK1γ1 interacted with wild-type, ST/A, and ST/E Claspin equally before and after heat stress (data not shown), Cdc7 bound to wild-type and ST/E Claspin, and much less to ST/A Claspin either with or without heat treatment (**Fig. 6B**, lanes 2-4 and 6-8), although the interaction was slightly stimulated after heat treatment. Furthermore, Cdc7 interacted with an aa897-1339 polypeptide containing the Cdc7-binding acidic patch (AP) domain when expressed in 293T cells. Significantly, Cdc7 bound to the ST/E mutant more efficiently than to wild-type, while it bound to the ST/A mutant less efficiently (**Supplementary Fig. S4**). These results suggest that C-terminal phosphorylation may stimulate Claspin interactions with Cdc7.

In **Figure 4A**, purified ST/E Claspin was phosphorylated even in the absence of added kinase. One possible explanation for this is that ST/E Claspin interacts readily with Cdc7, and that, during purification with anti-Flag agarose, Cdc7 may have been co-purified. To determine whether Cdc7 is responsible for phosphorylation of Claspin, we repeated the kinase assay in the presence of a specific Cdc7 inhibitor, and indeed, TAK-931 significantly reduced phosphorylation of ST/E Claspin (**Supplementary Fig. S5**, lane 4). These findings suggest that phosphorylation of Claspin’s C-terminal segment promotes Cdc7 recruitment.

### Mechanisms of heat-inducible Chk1 activation through C-terminal phosphorylation of Claspin

We hypothesize that the C-terminal domain of Claspin (amino acids 1319–1337: KHLKTDDSTSGLTRSIFKYL; theoretical pI: 10.04) interacts with AP, a domain in Claspin responsible for recruiting Cdc7. Phosphorylation of the C-terminal tail would decrease its interaction with AP, allowing AP to facilitate recruitment of Cdc7.

To explore this possibility, we used AlphaFold 3 (51) to predict the structure and interactions between Claspin (aa897-1011), containing the Chk1 binding domain (CKBD) + AP and wild-type, ST/A and ST/E Claspin C-terminal peptides (aa1320-1339). Predictions from AlphaFold 3 showed that the ST/A mutant C-terminus was embedded within Claspin 897-1011aa, while the wild-type polypeptide was partially embedded, and the ST/E mutant was not embedded (**Fig. 6C**). Thus, the C-terminal peptide has the potential to interact with AP, and this interaction could compete with Cdc7 binding. Phosphorylation of the C-terminal polypeptide inhibits its interaction with the AP, allowing Cdc7 access (**Fig. 6C**).

To test above model, we performed pull-down assays using Flag-tagged Claspin fragments (aa897-1100) containing AP and HA-tagged C-terminal polypeptide (aa1102-1339). We observed stronger binding to the ST/A mutant C-terminal fragment compared to the WT or ST/E fragments (**Fig. 6D**, lanes 1-3). Furthermore, coexpression of HRI suppressed the interaction of the WT but not ST/A C-terminal polypeptide with AP. GCN2 also showed similar effects (**Fig. 6D**, lanes 4-7). Consistent with the idea that phosphorylation of the C-terminal tail allows Cdc7 to bind to Claspin and phosphorylate the CKBD to activate checkpoint, we found that overexpression of HRI or GCN2 was sufficient to activate Chk1 at 37°C (**Supplementary Fig. S6**, lanes 2 and 3).

### Phosphorylation of the C-terminal tail of Claspin is involved in checkpoint activation by other stresses

Our data above show that phosphorylation of Claspin at its C-terminal tail by eIF2α kinases activates replication checkpoint to prevent heat-induced DNA damage and promote cell survival during heat stress. We next investigated whether other types of stress can similarly activate checkpoint through the C-terminal tail of Claspin. We used HCT116 Claspin-deficient cells, stably expressing either wild-type or ST/A mutant Claspin. These cells were treated with various stressors: hydroxyurea (HU), UV irradiation, heat, NaCl (osmotic stress), H2O2 (oxidative stress), high glucose (HG), lipopolysaccharide (LPS, bacterial toxin), and arsenate (Ar). As expected, the ST/A mutation decreased heat-induced Chk1 activation (to 52% of the wild-type levels). Furthermore, the ST/A mutation reduced Chk1 activation to 84% and 50% of the wild-type amounts upon high glucose and LPS stresses, respectively, suggesting that the cellular response to high glucose and bacterial toxins may also adopt the ISR pathway to activate checkpoint (**Supplementary Fig. S7**). Chk1 activation by other stressors was not significantly affected by ST/A mutation.

## Discussion

It was previously reported that heat stress activates Chk1 in a manner dependent on Claspin as well as other proteins including Rad9, Rad17 and TopBP1 (18,20). However, the precise mechanism of heat-induced activation of replication checkpoint has not been known. This study presents evidence indicating that heat stress activates the integrative stress response kinases, GCN2 and HRI, which phosphorylate the C-terminal tail of Claspin allowing Claspin to recruit Cdc7 and activate checkpoint. We identified five critical phosphorylation sites (T1322, S1325, T1326, S1327, and T1330) near the C-terminus of Claspin, which are essential for checkpoint activation and cell survival during heat shock. These findings reveal a crucial role for Claspin in protecting cells from heat-induced DNA damage by regulating the replication stress checkpoint, slowing down DNA replication fork movement, and permitting DNA repair.

### A novel role of ISR in replication checkpoint activation by heat stress

The results presented here show that cells undergo increased DNA damage in the absence of the Claspin C-terminal phosphorylation, presumably due to unchecked replication during heat stress. DNA replication is inhibited under heat stress, and part of this inhibition depends on checkpoint. In agreement with previous reports, we find that Claspin is important for normal replication fork progression. No significant effect of ST/A or ST/E mutation was observed for recovery of fork speed, suggesting that C-terminal phosphorylation does not affect Claspin’s role at the fork. Claspin depletion slows fork progression even in heat-stressed conditions, and expression of the non-phosphorylatable ST/A mutant Claspin increases fork speed significantly more than that of wild-type or the phosphomimic ST/E (**Fig. 2D**). Thus, after heat, fork rate could be reduced in a checkpoint-dependent manner.

ATR is required for full activation of Chk1 during heat stress. We previously showed that heat stress strongly inhibits DNA replication (26), suggesting that heat stress could induce double stranded DNA breaks (52), or changes in chromatin architecture (53) to activate ATR. While we are not clear of the precise functional interaction between ATR and C-terminal Claspin phosphorylation in inducing Chk1 activation, we note that ATR binds to wild-type, ST/E, and ST/A Claspin equally (**Supplementary Figure S3**), suggesting that ATR and C-terminal phosphorylation work in parallel to cooperatively activate Chk1. eIF2α kinases are primarily localized in cytoplasmic or organellar compartments. However, they could change their cellular localization under specific stress conditions. Indeed, PKR was reported to be translocated to the nuclei in response to radiation treatment in human lung cancer cells (54). The effects of heat treatment on the cellular localization of HIR and GCN2 will need to be examined.

We propose the following model for Chk1 activation (**Supplementary Fig. S8**). Upon heat stress, HRI and GCN2 phosphorylate the C-terminal serine/ threonine residues of Claspin. This induces a conformational change, causing dissociation of the C-terminal segment from the AP of Claspin. The unmasked AP domain is then free to recruit Cdc7. Recruited Cdc7 phosphorylates the CKBD of Claspin facilitating Chk1 binding. ATR, recruited to Claspin independent of the C-terminal phosphorylation, activates Claspin-bound Chk1 (55). However, we cannot exclude the possibility that the phosphorylated C-terminal tail of Claspin directly facilitates the recruitment of Cdc7 to Claspin, since the binding of Cdc7 to Claspin is compromised by ST/A mutation even in the absence of heat treatment (**Fig. 6B**).

### Replication checkpoint and the heat shock response pathway

HSF1 (Heat Shock Factor 1) is a transcription factor that plays a crucial role in the cellular response to heat stress (56). When cells are exposed to heat shock, HSF1 is activated and binds to specific DNA sequences known as heat shock elements (HSEs) in the promoters of heat shock protein genes, leading to the production of heat shock proteins (HSPs). HSPs function primarily as molecular chaperones, facilitating the proper folding of proteins, preventing protein aggregation, and aiding in the refolding or degradation of misfolded proteins (57).

Some heat shock proteins interact with components of the ISR and regulate the activities of stress-sensing kinases (eIF2α kinases). Activity of these kinases stabilizes ATR, ensuring its proper function in the cellular response to DNA damage and replication checkpoint activation.

Besides inducing the expression of HSPs, HSF1 also modulates the expression of ATF4 (Activating Transcription Factor 4), a key regulator of the ISR. ATF4 upregulates the expression of genes involved in protein folding and stress responses, including some HSPs. Thus, the ISR and heat shock response pathways function in concert to facilitate cell survival under conditions of heat stress, and can contribute to replication checkpoint activation.

### Stress-induced phosphorylation of Claspin as a potential novel target of cancer therapy

Our findings show that the roles of eIF2α kinases extend beyond their traditional roles in regulating protein synthesis. GCN2 and HRI not only phosphorylate eIF2α to modulate translation, but also directly phosphorylate Claspin to control DNA replication checkpoint, linking the integrated stress response (ISR) pathway to the replication stress response pathway. This connection underscores the broader role of eIF2α kinases in coordinating cellular responses to environmental stresses, ensuring both protein homeostasis and genomic stability.

The identification of this phosphorylation-mediated regulation has significant implications, particularly in the context of cancer research. Cancer cells constantly experience high levels of replication stress and rely on robust checkpoint mechanisms to survive. Targeting the Claspin phosphorylation pathway by eIF2α kinases could offer new therapeutic strategies to sensitize cancer cells to stress, potentially enhancing the efficacy of existing cancer treatments.

### Potential crosstalk through Claspin phosphorylation between Replication checkpoint and cellular responses to various biological stresses

Our data indicate that C-terminal phosphorylation of Claspin is also involved in cellular responses to LPS and high glucose. Future research will determine whether this phosphorylation mechanism is also used in response to other stresses, including oxidative stress, nutrient deprivation, hypoxia, ER stress, and unfolded protein response, and whether other eIF2α kinases, such as PERK and PKR, contribute to Claspin regulation under different conditions. In addition, understanding the precise mechanisms of how cancer cells protect themselves against various biological stresses may lead to the identification of additional therapeutic targets.

Overall, our study reveals a novel role of eIF2α kinase-mediated phosphorylation of Claspin in heat stress-induced replication checkpoint, pointing to crosstalk between the conserved cellular stress response pathway and replication checkpoint. This mechanism ensures the maintenance of genome stability under stress and offers a novel therapeutic potential for treating diseases like cancer, where stress response pathways are often dysregulated.

### Limitations of the study

We have shown that ISR plays a role in heat-stress induced replication checkpoint activation in cellular assays, and is required for cell survival after heat stress. However, it is not clear whether ISR is required for cell survival after heat stress in vivo. Does heat along with inhibition of Claspin or ISR induce genome instability or facilitate cancer formation in vivo? This should be examined by using animal models. It would be of interest whether other forms of stresses known to activate ISR have similar effects on cell survival in vivo when the ISR-Claspin pathway is compromized.

Claspin is a highly unstructured protein, and our results indicate the importance of intramolecular interactions in regulating its functions. These interactions are likely to be regulated by protein modification, including phosphorylation (You et al. submitted). The structural dynamics of Claspin under various stresses or during cell cycle needs to be elucidated by high-speed AFM or visualization of the protein under CryoEM.

## Materials and Methods

### Cell lines

293T, HCT116, and U2OS cells were obtained from ATCC. Claspin flox/-Mouse Embryonic Fibroblasts (MEFs) were established from E12.5 embryos (11). HCT116-Claspin-myc-AID cells, stably expressing wild-type Flag, ST/A-Flag, or ST/E-Flag mutant Claspin, were generated by infection with recombinant lentiviruses expressing these cDNAs. Cells were cultured in Dulbecco’s Modified Eagle’s Medium (high glucose), supplemented with 15% fetal bovine serum, 2 mM L-glutamine, 1% sodium pyruvate, 100 U/ml penicillin, and 100 μg/ml streptomycin. To select for HCT116-Claspin-myc-AID cells stably expressing exogenous Claspin, 10 μg/ml Blasticidin was also added. Cells were maintained in a humidified atmosphere with 5% CO2 and 95% air at 37°C.

### Plasmid construction

DNA fragments encoding portions of Claspin or its mutant forms were obtained through PCR amplification to express truncated or mutant versions of Claspin. For the expression of Claspin mutants ST/A and ST/D, in which all serines and threonines within the 9 aa of 1322-1330aa were replaced with alanine or aspartic acid, respectively, was synthesized. Both wild-type and mutant Claspin fragments were inserted into the CSII-EF MCS-6His-Claspin-3Flag-P2A-Bsd (Blasticidin) vector (58) using the In-Fusion cloning method to construct lentiviral expression vectors for full-length Claspin WT, ST/A, or ST/D mutants.

### Antibodies, proteins, siRNA and peptide

The antibodies used in this study are as follows. Anti-human Claspin was developed in rabbit against a recombinant protein containing amino acids 896-1014 of human Claspin produced in *Escherichia coli* and sera was used at a dilution of 1:1000. Anti-mouse Claspin was developed in rabbit against the polypeptide LKTNGSSPGPKRSIFRYLES (amino acids 1296-1315 of mouse Claspin), and the sera was used at a dilution of 1:500. Anti-Tubulin (sc23948, 1:5000) antibody was from Santa Cruz Biotechnology. Anti-Chk1 (K0086-3, 1:500), anti-DDDDK-tag (M185-3L, 1:1000), and anti-Cdc7 (K0070-3, 1:1000) antibodies were obtained from MBL. Anti-Rat IgG Alexa Fluor 555 (405420, 1:300) and FITC-anti-BrdU (364104) were obtained from Biolegend. Anti-Mouse IgG Alexa Fluor 488 (A-11017, 1:100) was obtained from Invitrogen. Anti-BrdU clone B44 (#347580, 1:5) was obtained from BD. Anti-BrdU (ab6326, 1:30) was obtained from Abcam. Anti-Chk1 S345 (#2341, 1:1000) antibodies were obtained from Cell Signaling. RPA32 phospho-S4/S8 (A300-245A, 1:1000) and anti-CK1γ1 (orb37898, 1:1000) antibodies were from Bethyl and Biorbyt, respectively. The proteins and peptide used in this study are as follows. Purified Chk1 (02–117), HRI (05–154), and GCN2 (05–153) kinases were obtained from Carna Bioscience. CHKtide (C10-58) peptide was obtained from SIGNALCHEM BIOTECH INC. λ phosphatase (PPase) was from NEB.

siRNAs for mouse GCN2 and HRI were as follows. mus GCN2 Sense 5’: AGUUGGUGACGAAAGAAAU-UU; mus GCN2 Anti-Sense 3’: UU-UCAACCACUGCUUUCUUUA. mus HRI Sense 5’: AGCAGUUCGUCCAUUGUCU-UU; mus HRI Anti-Sense 3’: UU-UCGUCAAGCAGGUAACAGA.

### Expression of recombinant proteins in 293T cells

1.6 μg of expression plasmid DNA in 100 μL of 150 mM NaCl was mixed with 100 μL of 150 mM NaCl supplemented with 7 μL of 1 mg/mL PEI (polyethylenimine ’MAX’ [MW 25,000; Cat. 24765; Polyscience, Inc.]). After a 30-min incubation at room temperature, the solution was added to 293T cells cultured in six-well plates, each containing 2 mL of fresh DMEM. For 15 cm plates, the amounts of DNA/ reagents were increased by 10-fold.

### Expression of proteins using lentivirus vectors

293T cells were transfected with the lentivirus vector expressing wild type or mutant Claspin (on CSII-EF MCS-6His-Claspin-3Flag-P2A-Bsd), pMDLg/pRRE, pRSV-Rev and pMD2.G. Virus-containing medium were harvested at 2 days (for lentivirus) after transfection and the viruses were concentrated by centrifugation. The concentrated Claspin-expressing lentiviruses with 4 μg/ml polybrene were used for infection of HCT116-Classpin-AID-myc. At 2 days after infection, 10 μg/ml Blasticidin was added and resistant cells were selected for 5 days.

### Protein purification

293T cells of five 15 cm dishes were incubated for 40 hrs after transfection, then harvested and lysed as previously described (58). The proteins bound to ANTI-FLAG® M2 Affinity Gel (Sigma-Aldrich; A2220) were recovered from the supernatants and washed with Flag wash buffer [50 mM NaPi (pH 7.5), 10% glycerol, 300 mM NaCl, 0.2 mM PMSF, and protease inhibitor tablet]. The bound proteins were then eluted using Flag elution buffer [50 mM NaPi (pH 7.5), 10% glycerol, 100 mM NaCl, 200 μg/ml 3xFlag peptide (Sigma), 0.1 mM PMSF, and protease inhibitor tablet].

### Western blotting

Proteins in SDS-sample buffer were incubated at 96°C for 3 min, separated by SDS-PAGE, and then transferred onto Hybond ECL membranes (GE Healthcare). Western blot analysis was performed using the specified primary antibodies. The blots were then incubated for 1 hr with a horseradish peroxidase-conjugated secondary antibody. Protein detection was carried out using Chemi-Lumi One Super Western Blotting Substrate (Nacalai), and images were captured using the LAS3000 system (Fujifilm).

### Mass Spectrometry Analysis of Claspin

293T cells stably expressing Claspin-10xFlag (15x15 cm dishes) were cultured at either 37°C or 42°C for 2 hrs and then harvested. The harvested cells were lysed using RIPA buffer (50 mM Tris-HCl [pH 8.0], 150 mM NaCl, 0.5 % w/v sodium deoxycholate, 1.0 % NP-40, and 0.1 % SDS) supplemented with 1 mM DTT, 1 mM Na₃VO₄, 50 mM NaF, 0.1 mM ATP, 0.5 mM PMSF, and a protease inhibitor tablet. The supernatants were incubated with ANTI-FLAG® M2 Affinity Gel at 4°C for 1 hr. After washing with PBS containing 0.01 % Tween-20, the bound proteins were separated on a 4–20 % gradient gel and stained using Coomassie Brilliant Blue (CBB). The Claspin protein was excised from the gel, digested with trypsin, and phospho-threonines or phospho-serines were analyzed by mass spectrometry.

### Cell cycle analysis

U2OS cells were treated with siRNA for 4 hrs (by using oligofectamine reagent), followed by the addition of 2.5 mM thymidine and incubation at 37°C for 18 hrs. Cells were then washed twice with pre-warmed serum-free medium or PBS. Fresh pre-warmed medium was added, and cells were released at 37°C for 9 hrs. Next, 25 ng/ml nocodazole was added, and cells were incubated at 37°C for 12 hrs. Cells were gently resuspended in pre-warmed serum-free medium, centrifuged, and the supernatant was discarded. Cells were washed twice with pre-warmed serum-free medium, followed by the addition of fresh pre-warmed medium. Cells were then released at 37°C or 42°C. At the indicated time points, cells were harvested and fixed in 75% ethanol at −20°C. After washing with wash buffer (0.5% bovine serum albumin in phosphate-buffered saline), cells were incubated with 25 μg/ml propidium iodide and 100 μg/ml RNase A for 30 min at room temperature. Finally, cell cycle distribution was analyzed using fluorescence-activated cell sorter (FACS).

### Colony formation assays

HCT116 Claspin-myc-AID cells and the same cells stably expressing either wild-type or mutant Claspin were treated with or without 0.5 mM auxin for 3 hrs, and were cultured further at 37°C or 42°C for 90 min. One thousand cells were plated in 6-well plates with or without 0.5 mM auxin and allowed to form colonies for 3 weeks at 37°C. HCT116 Claspin-myc-AID cells treated with or without 0.5 mM auxin and 10 nM GCN2 inhibitor (GCN2-IN-1(A-92)) for 3 hrs were cultured at 37°C or 42°C for 90 min. Five hundred cells were plated in 6-well plates with or without 0.5 mM auxin and allowed to form colonies for 3 weeks at 37°C.

In both cases, following colony formation, cells were washed twice with 2 mL of 1X PBS, fixed with methanol at room temperature for 20 min, and stained with Coomassie Brilliant Blue at room temperature for 1 hr. The plates were then washed with water, allowed to dry, and the number of colonies was counted.

### DNA fiber

HCT116 Claspin-myc-AID cells and the same cells stably expressing either wild-type or mutant Claspin were treated with 0.5 mM auxin for 3 hrs, and were cultured further at 37°C or 42°C for 90 min. The cells were pulse-labeled with 25 μM CldU (Sigma; C6891) for 15 min, followed by 250 μM IdU (TCI; I0258) for an additional 15 min. Labeled cells were then placed on glass slides, and DNA fibers were spread in a lysis buffer containing 0.5% SDS, 200 mM Tris (pH 7.4), and 50 mM EDTA. After drying, the DNA was fixed using Carnoy solution (methanol: acetic acid = 3:1) for 3 min. DNA was denatured with 2.5 N HCl for 30 min and then neutralized with 0.1 M sodium borate for 3 min. For immunodetection of labeled tracks, fibers were incubated with primary antibodies, rat anti-BrdU (for CldU; Abcam, ab6326) and mouse anti-BrdU (for IdU; BD Biosciences, 347580), for 1 hr at room temperature. The fibers were then detected with corresponding secondary antibodies, anti-mouse Alexa 488 (Abcam; ab150113) and anti-rat Alexa 555 (Abcam; ab150158), for 1 hr. Slides were examined on the all-in-one fluorescence microscope (KEYENCE; BZ-X800). In each assay, more than 66 tracks were measured to estimate fork rate.

### Comet assays

2 × 10^4^ HCT116 Claspin-myc-AID cells as well as HCT116 Claspin-myc-AID cells stably expressing either wild-type or mutant Claspin, treated with or without 0.5 mM auxin for 3 hrs, were cultured at 37°C or 42°C for 90 min. The comet assay was performed using the Comet Assay Kit (Abcam; ab238544) following the provided protocol. Briefly, harvested cells were resuspended at 1 × 10^5^ cells/mL in ice-cold PBS. 8 μL of the cell suspension was mixed with 80 μl of 1% low melting point agarose and transferred onto the top of a coated agarose base layer on a slide. The slide was then placed at 4°C for 20 min. The slide was immersed in pre-chilled lysis buffer (1X lysis solution; 100 mM EDTA; 2.5 M NaCl; 10% DMSO) for 60 min at 4°C. It was then transferred to a pre-chilled alkaline solution (1 mM EDTA; 0.3 M NaOH) for 30 min at 4°C. Afterward, the slide was washed twice in pre-chilled TBE buffer for 5 min each and placed in an electrophoresis chamber for 15 min at 25 volts. The slide was transferred to pre-chilled distilled water for 2 min, three times, and then replaced with cold 70% ethanol for 5 min before being air-dried. After staining with DAPI (1 μg/mL) for 15 min, the slides were examined on the all-in-one fluorescence microscope with a DAPI filter (KEYENCE; BZ-X800). Comet tail length was measured using the BZ-X Analyzer software.

### *In vitro* kinase assays

Approximately 100 ng of purified wild-type Chk1 or kinase-dead Chk1, and 200 ng of CHKtide peptides (when applicable), were incubated with or without GCN2 or HRI (20 ng) in kinase reaction buffer (40 mM Hepes-KOH [pH 7.6], 2.5 mM spermine, 5 mM MgCl₂, 0.5 mM EGTA, 0.5 mM EDTA, 1 mM Na₃VO₄, 1 mM NaF, 2 mM DTT, 10 μM ATP, and 1 μCi [γ-³²P]ATP) for 1 hr at 37°C. Similarly, λPPase-treated purified wild-type or mutant full-length Claspin, or wild-type or mutant Claspin polypeptides, which were pre-treated with PhosSTOP™ (Sigma-Aldrich; 4906845001) at 2°C for 1 hr, were incubated with or without GCN2 or HRI (20 ng) in the same kinase reaction buffer under the same conditions. After incubation, one-fourth volume of 5x sample buffer was added, and the mixtures were heated at 96°C for 3 min. The samples were analyzed by SDS-PAGE, followed by Coomassie Brilliant Blue (CBB) staining and autoradiography.

### Quantification and statistical analysis

Error bars represent the mean ± standard deviation (s.d.) calculated from three independent replicate experiments. p-values were determined using a Student’s two-tailed t-test.

## Acknowledgements

We thank Naoko Kakusho and Rino Fukatsu for exelent technical assistance. This work was supported by JSPS KAKENHI (Grant-in-Aid for Scientific Research (A) [Grant Numbers 20K21410 and 20H00463 (to H.M.)]; Hirose international scholarship foundation (to C-C.Y.), and JSPS KAKENHI grant number JP23K06674 (HG). We thank Dr. Hiroyuki Sasanuma for continuous support. We thank Masato Kanemaki (National Institute of Genetics) for providing IAA-inducible protein degradation system. We would like to thank all the members of the laboratory for helpful discussion.

## Author contributions

C-C.Y. performed most of the experiments and contributed to the analysis of the results described in the paper. H.M. performed some of the experiments (plasmid construction, HU time course etc.) experiments. H.G. constructed HCT116 Claspin-mAID cell lines.

S.O. constructed MEF 4KO and its derivatives. C-C.Y. and H.M. designed the experiments and wrote the manuscript with input from all authors. H.M. obtained fundings and has overall responsibility for the study.

## Legends to Supplementary Figures

**Supplementary Figure S1. Phosphorylation of the C-terminal of Claspin under heat shock stress.**

Mass spectrometry analysis of 293T cells stably expressing Claspin-10xFlag. Cells were cultured in 15 x 15 cm plates at 42°C and harvested at 0 and 2 hrs. Samples were analyzed by SDS-PAGE followed by Coomassie Brilliant Blue (CBB) staining, and mass spectrometry was performed. Residues in red represent amino acids phosphorylated specifically at 2 hrs but not at 0 hr.

Comparison of the C-terminal sequences of Claspin homologs across various species. Clusters of serine/ threonine residues are highlighted in bold.

**Supplementary Figure S2. Heat-induced activation of Chk1 depends on Claspin and its C-terminal phosphorylation.**

HCT116 Claspin-myc-AID cells stably expressing either wild-type (WT) or ST/A mutant Claspin were treated with 0.5 mM auxin for 3 hrs to induce Claspin degradation. Cells were then incubated at 37°C or 42°C for 90 min. The whole-cell extracts were analyzed by western blotting with the indicated antibodies.

**Supplementary Figure S3. Claspin interacts with ATR in a manner independent of the C-terminal phosphorylation.**

The wild-type (WT), ST/A or ST/E mutant Claspin were overexpressed in 293T cells. Cell lysates were immunoprecipiated with anti-Flag (M2) beads, and co-immunoprecipitation of ATR was examined.

**Supplementary Figure S4. The Claspin ST/E mutant strongly binds to Cdc7 after transient expression in mammalian cells.**

293T cells were transfected with plasmids expressing Flag-tagged wild-type (WT), ST/A, or ST/E Claspin C-terminal polypeptide (aa897-1339). Cell lysates were immunoprecipitated with anti-Flag (M2) beads, and co-immunoprecipitation of Cdc7 was examined.

**Supplementary Figure S5. The ST/E Claspin purified from mammalian cells may contain Cdc7 protein.**

Purified wild-type and ST/E mutant Claspin proteins were incubated in the phosphorylation reaction condition with or without 20 μM TAK-931 (Cdc7 kinase inhibitor) at 30°C for 30 min. The reaction mixtures were subjected to SDS-PAGE. The gel was stained with Coomassie Brilliant Blue (CBB) (left panel) and then autoradiographed (right panel).

**Supplementary Figure S6. Activation of Chk1 by eIF2α kinases.**

HRI or GCN2 was overexpressed in 293FT cells and cultured at either 37°C or 42°C for 90 min. The whole-cell extracts were analyzed by western blotting using the specified antibodies to assess Chk1 activation.

**Supplementary Figure S7. Potential roles of the C-terminal phosphorylation of Claspin on Chk1 activation by various stresses.**

HCT116 Claspin-myc-AID cells stably expressing wild-type (WT) or ST/A mutant Claspin were treated with auxin for 3 hrs and then exposed to the indicated stressors for 2 hrs. Chk1 activation was examined in the whole cell extracts. The numbers below the pChk1 panel indicate the relative intensity of pChk1 bands (normalized by the level of Chk1 protein) in ST/A samples compared to the WT samples. Conditions of the stresses were as follows. Heat, 42°C; NaCl, 50 mM; H_2_O_2_, 50 µM; HG (high glucose), 30 mM Glucose; LPS, 2 µg.ml; Ar (arsenate), 400 µM.

**Supplementary Figure S8. A model on Claspin-dependent Chk1 activation by heat stress.**

The C-terminal region of Claspin normally interacts with and masks the AP domain. Upon heat stress, ISR kinases phosphorylate the C-terminal region of Claspin. This phosphorylation event causes the C-terminus to dissociate from AP, promoting Cdc7 binding to AP and activating Chk1. ATR is recruited to Claspin and may activate Chk1 bound to Chk1 in a pathway separate from the ISR kinase-Cdc7-Chk1 pathway.

