## Supplementary Figures for "Phosphorylation of Claspin by elF2α kinase protects cells from heat stress"

## (A)

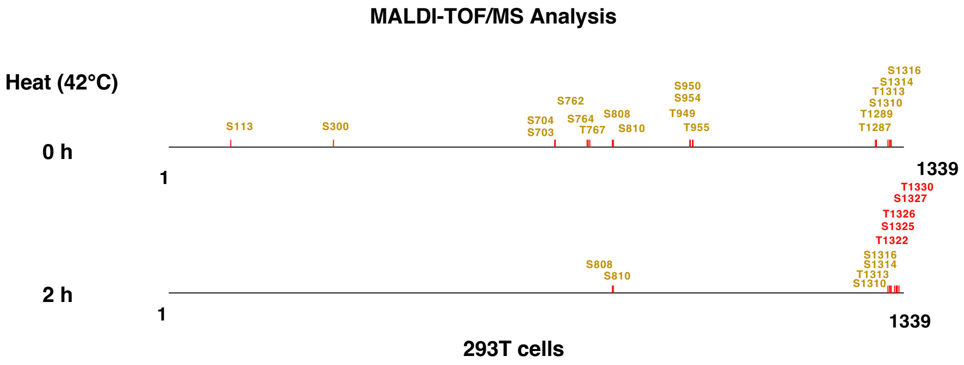


## (B)

| | | |
|---|---|---|
| **Claspin Homo sapiens** | **T**DD**STS**GL**T**RSIFKYLE-S | **1339** |
| **Claspin Bos taurus** | IDD**STS**E**S**KQSIFKYLE-S | **1327** |
| **Claspin Mus musculus** | **T**NG**SS**PGPKRSIFKYLE-S | **1315** |
| **Claspin Xenopus laevis** | RD**ST**P**T**VK**S**RSIFQLLE-- | **1285** |
| **Mrc1 Schizosaccharomyces** | Q**S**ANPPRLLASLNNYSDFD | **1019** |

**Yang et al. Supplementary Fig. S1**

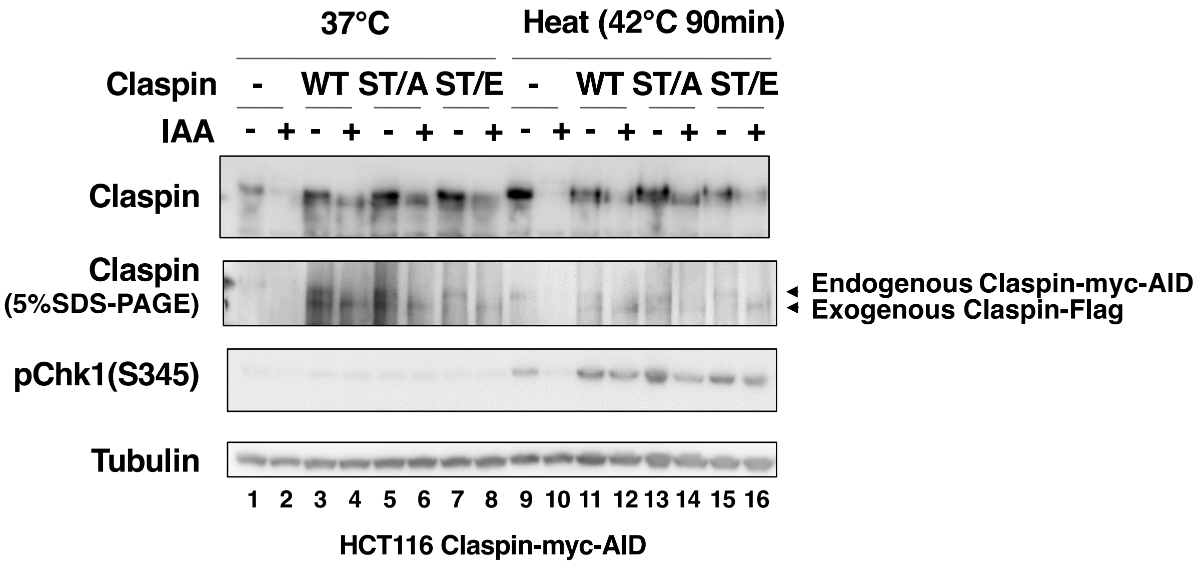


Yang et al. Supplementary Fig. S2

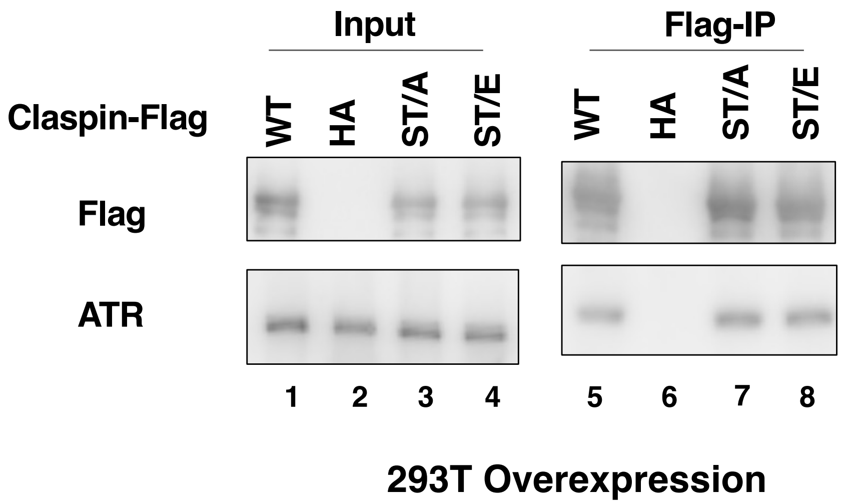


Yang et al. Supplementary Fig. S3

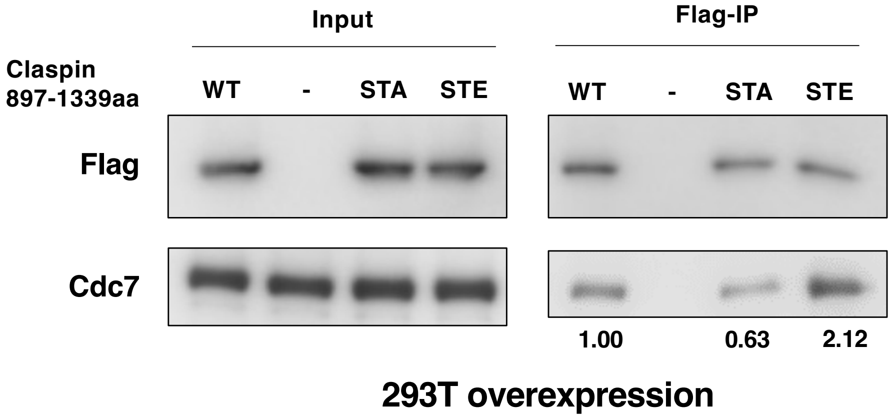


Yang et al. Supplementary Fig. S4

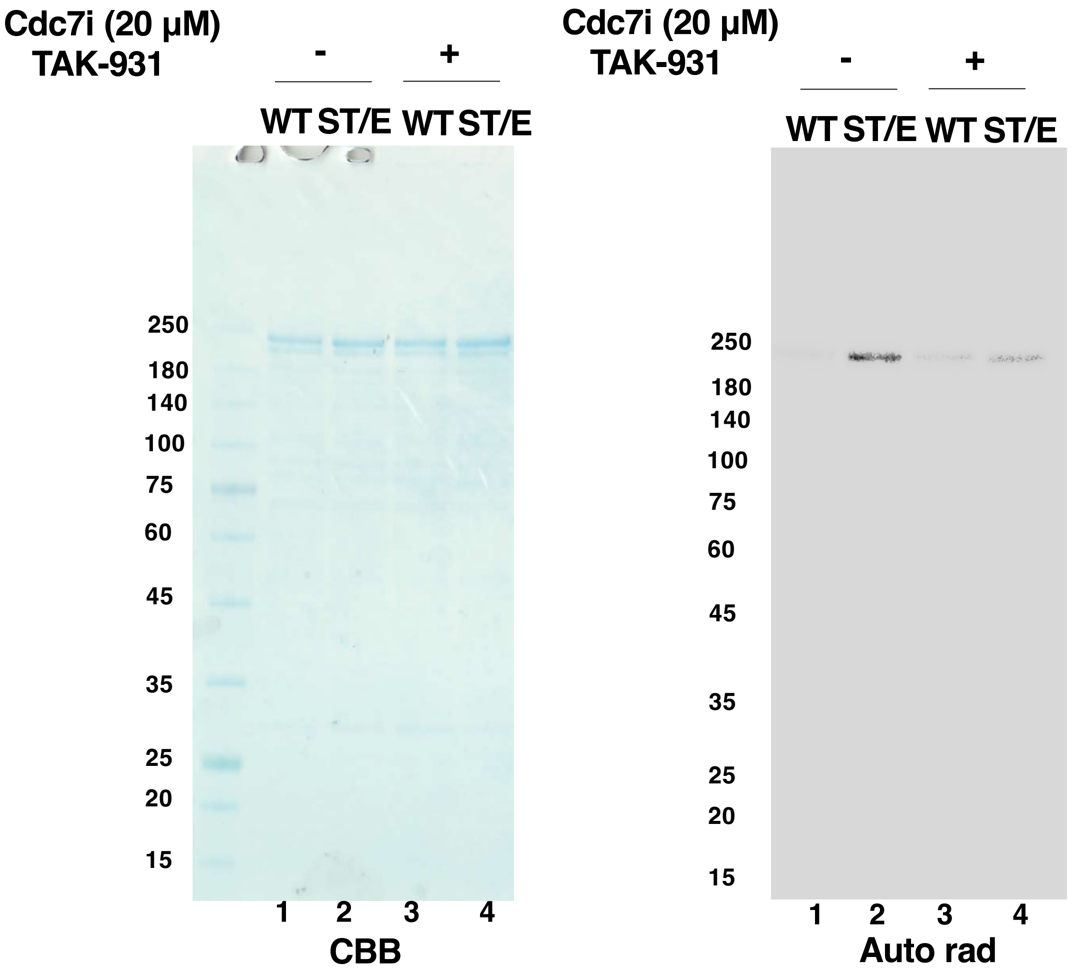


Yang et al. Supplementary Fig. S5

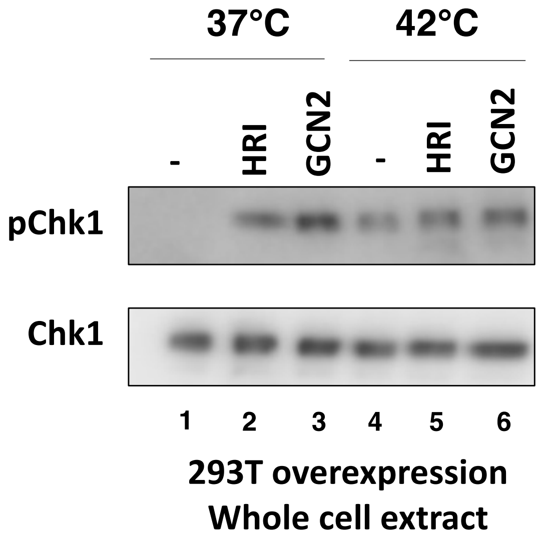


Yang et al. Supplementary Fig. S6

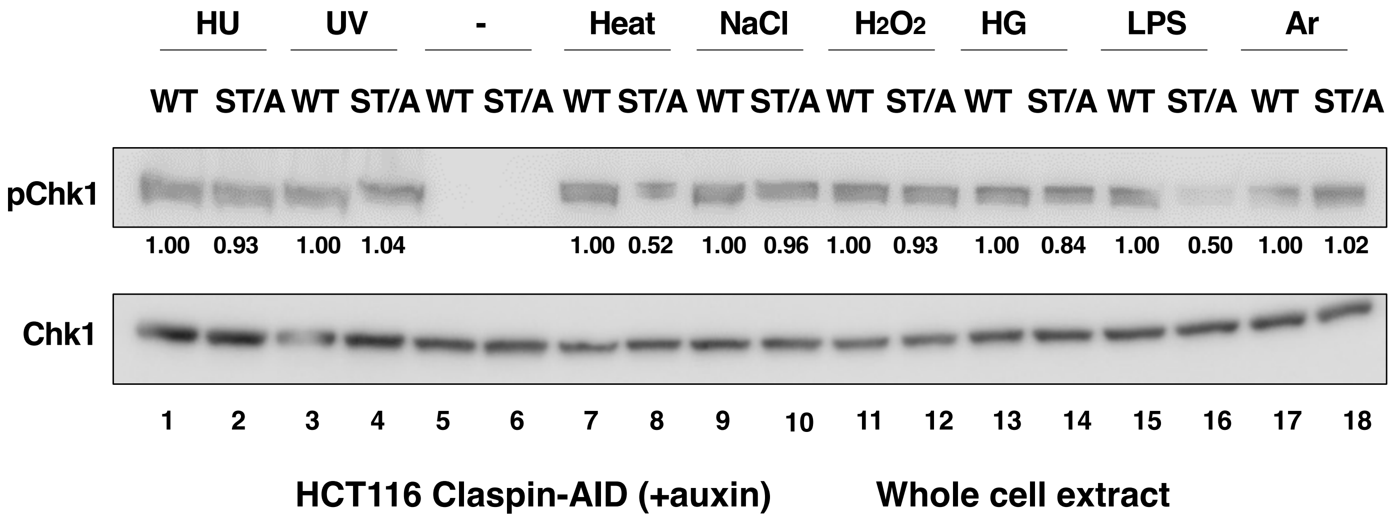


Yang et al. Supplementary Fig. S7

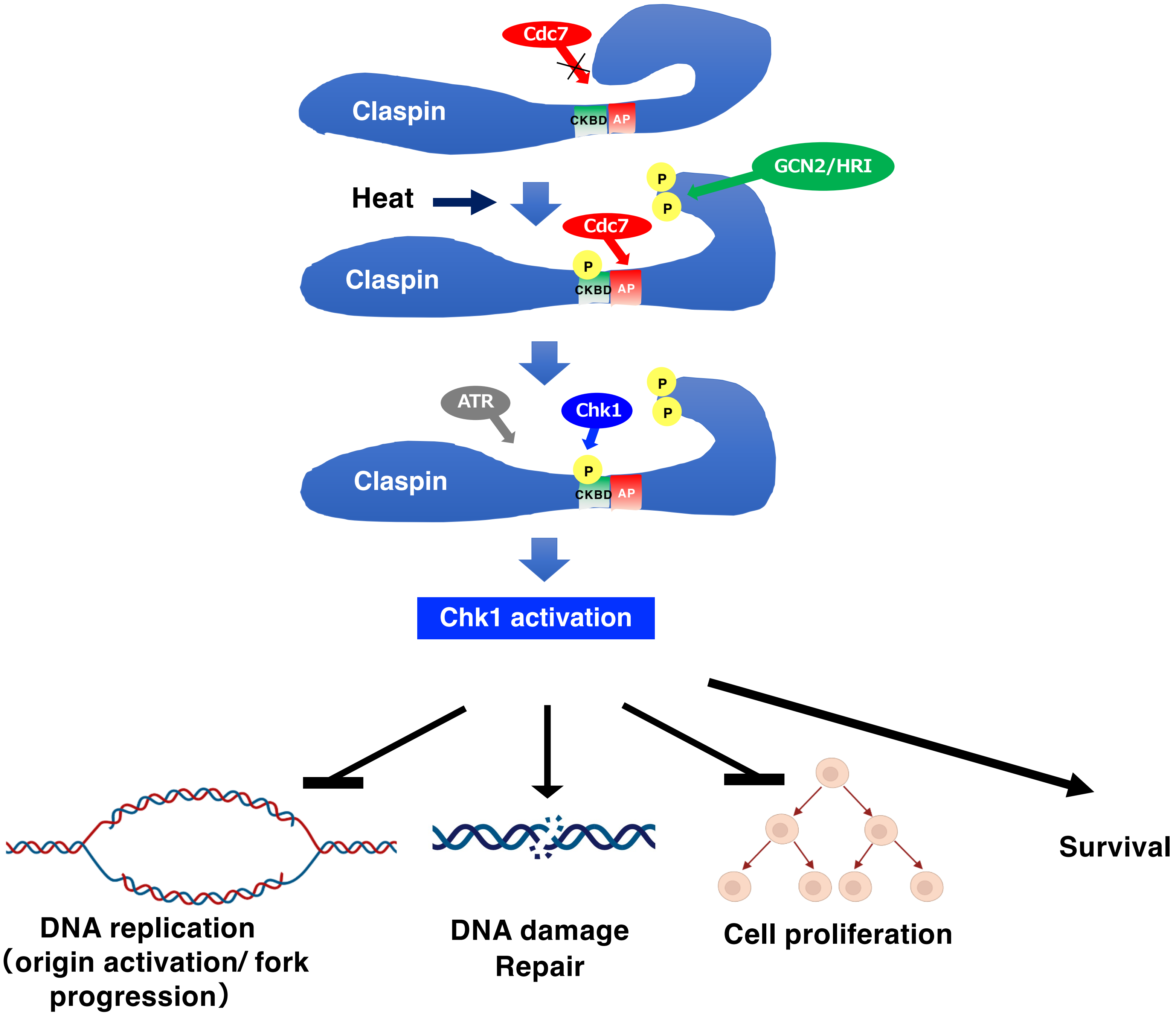


Yang et al. Supplementary Fig. S8
